# Predicting Phylogenetic Bootstrap Values via Machine Learning

**DOI:** 10.1101/2024.03.04.583288

**Authors:** Julius Wiegert, Dimitri Höhler, Julia Haag, Alexandros Stamatakis

**Affiliations:** Computational Molecular Evolution Group, Heidelberg Institute for Theoretical Studies, Heidelberg, Germany; Biodiversity Computing Group, Institute of Computer Science, Foundation for Research and Technology - Hellas, Heraklion, Crete, Greece; Institute for Theoretical Informatics, Karlsruhe Institute of Technology, Karlsruhe, Germany

## Abstract

**Summary:** Estimating the statistical robustness of the inferred tree(s) constitutes an integral part of most phylogenetic analyses. Commonly, one computes and assigns a branch support value to each inner branch of the inferred phylogeny. The most widely used method for calculating branch support on trees inferred under Maximum Likelihood (ML) is the Standard, non-parametric Felsenstein Bootstrap Support (SBS). Due to the high computational cost of the SBS, a plethora of methods has been developed to approximate it, for instance, via the Rapid Bootstrap (RB) algorithm. There have also been attempts to devise faster, alternative support measures, such as the SH-aLRT (Shimodaira–Hasegawa-like approximate Likelihood Ratio Test) or the UltraFast Bootstrap 2 (UFBoot2) method. Those faster alternatives exhibit some limitations, such as the need to assess model violations (UFBoot2) or meaningless low branch support intervals (SH-aLRT). Here, we present the Educated Bootstrap Guesser (EBG), a machine learning-based tool that predicts SBS branch support values for a given input phylogeny. EBG is on average 9.4 (*σ* = 5.5) times faster than UFBoot2. EBG-based SBS estimates exhibit a median absolute error of 5 when predicting SBS values between 0 and 100. Furthermore, EBG also provides uncertainty measures for all per-branch SBS predictions and thereby allows for a more rigorous and careful interpretation. EBG can predict SBS support values on a phylogeny comprising 1654 SARS-CoV2 genome sequences within 3 hours on a mid-class laptop. EBG is available under GNU GPL3.

**Data and Code Availability:** github.com/wiegertj/EBG

github.com/wiegertj/EBG-train

## 1 Introduction

Inferring phylogenetic trees under the Maximum Likelihood (ML) criterion is time- and resource-intensive, as the number of possible tree topologies increases super-exponentially with the number of taxa under study. As a consequence, tree search algorithms, as implemented in RAxML-NG [18], conduct their search for a best-known, yet not necessarily globally optimal, tree topology via a plethora of distinct heuristics. As there is no guarantee that the tree search will converge to the *globally* optimal tree, subsequent analyses to quantify the uncertainty are necessary. Such uncertainty analyses constitute an integral as well as routine component of current phylogenetic analysis pipelines [15]. The most common approach to quantify uncertainty is to infer various flavors of inner branch support values. The standard technique to calculate branch support values on a resulting phylogeny under ML is the Standard, non-parametric Felsenstein Bootstrap Support (SBS) [7]. The SBS randomly samples the alignment sites of the Multiple Sequence Alignment (MSA) with replacement to create a set of replicate MSAs (called bootstrap replicates). On each such replicate, one then infers a respective ML tree. Thus, the SBS yields a set of bootstrap replicate ML trees. This SBS procedure is time- and resource-consuming, due to the high computational cost of conducting a phylogenetic tree inference on each replicate. Note that typically 100-500 replicate trees need to be inferred to obtain stable support values [23].

To alleviate this computational bottleneck, a plethora of alternative, faster methods to infer branch support have been proposed. For example, Stamatakis et al. [29] propose the Rapid Bootstrap (RB) as part of the phylogenetic inference tool RAxML [28] as a faster alternative to the SBS. RB uses a heuristic approach that implements a more superficial ML tree search to approximate the SBS values and reduce computational costs. On multiple large datasets Stamatakis et al. [29] show, that RB support values are highly correlated with the SBS values (Pearson correlation between 0.92 and 0.99). The Ultrafast Bootstrap UFBoot [21] and its current version UFBoot2 [14] interweave parametric (substitution model-dependent) and non-parametric aspects. While the tree space sampling of UFBoot2 is parametric, it deploys a non-parametric bootstrap sampling of the MSA. UFBoot2 yields easy-to-interpret, unbiased branch support values, the UltraFast Bootstrap Support (UFBS). According to the authors’ experiments, UFBoot2 is extremely fast, as, on median, it is 778 times faster than SBS and 8.4 times faster than RB.

Both, RB (optionally) and UFBoot2 employ an iterative approximation of their branch supports until they either meet a stopping criterion or reach a maximum number of iterations. Thus, the runtimes of these methods can vary, since they depend on the input MSA, which determines if and when the support value calculations will converge.

Anisimova and Gascuel [3] propose an alternative definition of branch support based on a parametric method for branch support estimation: The approximate likelihood ratio test (aLRT). The aLRT compares the two best NearestNeighbor-Interchange (NNI) moves at each inner branch via an LRT test, to calculate a branch support value. To accelerate the NNI likelihood evaluation, aLRT only optimizes the branches adjacent to the branch of interest. As aLRT is parametric, it can be sensitive to substitution model violations. Substitution model violations occur, for example, when we choose a substitution model that is too simple for the data at hand. To correct for those violations, Guindon et al. [10] propose the Shimodaira–Hasegawa-like (SH-like) aLRT, which constitutes a non-parametric version of the aLRT. The aLRT and SH-like aLRT only focus on local perturbations of the given ML topology for which they calculate supports. This can induce overconfidence in branches if there exist other highly likely, yet topologically substantially distinct tree topologies [10]. One recent example of such a tree space with a multitude of topologically highly distinct yet almost equally likely tree topologies is a phylogeny of SARS-CoV2 genome sequences [22].

Despite the availability of these tools, SBS remains an important approach for measuring branch support [6, 1, 20]. Guindon et al. [10] propose to combine the SBS with the SH-like aLRT to obtain a holistic estimate of branch robustness. UFBoot2 is less vulnerable to severe model violations than UFBoot [14]. Yet UFBoot2 still requires an additional step to assess such violations. SBS is inherently robust against model violations, as it is non-parametric.

Here, we present the Educated Bootstrap Guesser (EBG), a machine learningbased approach for predicting SBS values. Predicting SBS values on 220 empirical MSAs with EBG is on average 9.4 (*σ* = 5.5) times faster with respect to time-to-completion compared to UFBoot2. Based on these experiments, we show that EBG can also predict whether a specific branch will exceed a specific SBS threshold (e.g., *t* := 80) in a 1000 replicate SBS run with a balanced accuracy of at least 0.91.

## 2 Materials and Methods

### 2.1 Phylogenetic Bootstrap

On each replicate, the SBS infers a corresponding ML tree, which yields a set of replicate trees. We can use those trees to either construct a consensus tree or to map confidence values onto a reference tree (typically the best-known ML tree) as SBS values. The SBS value for a specific inner branch (also referred to as non-trivial split or bipartition) is the frequency of occurrence of this branch in the replicate tree set. Common representations of the SBS values are percentage values between 0 and 100, or fractions between 0 and 1. In the following, we represent SBS values as percentage values between 0 and 100.

### 2.2 Problem Formulation

We address the challenge of predicting the SBS via two distinct steps: regression and classification. The EBG regressor directly predicts the respective SBS values for *all* inner branches of a given ML tree. In the classification approach, EBG then predicts the probability of the SBS value of each single inner branch to exceed a given SBS threshold using the regressor output as input. This classification step is based on the SBS interpretation by Felsenstein and Kishino [8]. The authors propose to interpret the quantity of one minus the SBS value as a *p*-value for the null hypothesis of the branch *not* forming part of the true tree. However, Susko [30] shows that one minus the SBS is too conservative as a *p*-value. Given an SBS value *>* 95, the probability of the branch not being in the true tree is substantially below 5%. Their experiments suggest that, depending on the characteristics of the true tree, an SBS between 70 and 85 yields a 5% false positive bound. Consequently, we use this SBS range for our classification step. More specifically, we predict the probability for each single branch *i* to exceed a specific SBS threshold *t*, that is, *SBS*_*i*_ *> t* with *t ∈* {70, 75, 80, 85}.

With our novel EBG tool, we address both, the regression and the classification challenge. In addition, we estimate the uncertainty of the resulting predictions.

### 2.3 Training Data

Trost et al. [32] demonstrate, that machine learning algorithms can easily distinguish between simulated and empirical MSAs with high accuracy and conclude that sequence simulations do not fully capture all characteristics of empirical MSAs. Consequently, we exclusively use empirical MSAs to train EBG. We obtained the empirical MSAs from TreeBASE [25]. As TreeBASE only contains MSAs of published studies, we assume that they are representative of commonly analyzed datasets by practitioners.

We used 1496 MSAs (containing DNA and Amino Acid (AA) sequences) for training and evaluating EBG. In addition, we randomly selected 220 additional MSAs for our final comparison with RB, UFBoot2, and SH-like aLRT. We noticed that approximately half of the MSAs in TreeBASE contain at least two exactly identical sequences, and therefore decided to remove all duplicate sequences before training EBG. We selected the MSAs based on the Pythia difficulty [12]. Pythia quantifies the difficulty of phylogenetic analysis under the ML criterion. To obtain a comprehensive representation of the Pythia difficulty spectrum in ML tree inference, we therefore predicted the Pythia difficulty for each MSA in TreeBASE using Pythia. Subsequently, we selected MSAs representing distinct Pythia difficulty levels to cover both, easy, and challenging MSAs.

For each MSA, we inferred 100 ML trees under the GTR+G substitution model using RAxML-NG [18]. To obtain SBS “ground truth” values as a training target for EBG, we performed one SBS run with 1000 replicates for each MSA using RAxML-NG. Our final training dataset comprises approximately 80 000 inner branches and their corresponding SBS value.

### 2.4 Feature Engineering

To predict the SBS values, we computed a plethora of MSA features but also features based on the respective best-known ML tree, including the model parameter estimates. EBG uses a total of 23 features for the prediction (Table 1).

**Table 1:** Overview of the features subset used in EBG.

| *Feature* |
| --- |
| Parsimony bootstrap support (PBS) |
| Parsimony support (PS) |
| Normalized ML branch length |
| # child inner branches |
| Skewness PBS |
| Mean Robinson-Foulds distance PB |
| Mean parsimony mutation per side (PMS) |
| ML Branch length |
| Max. PSF |
| Coefficient of variation PMS |
| Max. PBS children* |
| Mean PBS parents |
| Max. PS children* |
| Branch number (ordered by level-order traverse) |
| Skewness PMS |
| Branch length ratio bipartition |
| Max. PS children* |
| Min. PS children |
| Std. PBS parent branches |
| Std. PBS child branches |
| Mean closeness centrality bipartition ratio |
| Min. PS children* |
| Min. PBS children* |
| *: weighted by branch length |

The majority of features are extracted from a set of parsimony starting trees (henceforth simply referred to as parsimony trees) we inferred using RAxML-NG. RAxML-NG infers parsimony starting trees via a randomized stepwise addition order algorithm (−−start-option). The development of Pythia showed that by using the computationally substantially less expensive parsimony trees, we can accurately predict features of the ML tree space [12]. Due to the high prediction accuracy of Pythia, we therefore expect that parsimony-based features will also be useful for predicting SBS values.

We calculated a set of 12 features based on parsimony trees. Those 12 features are subdivided into Parsimony Support (PS) and Parsimony Bootstrap Support (PBS) features. Figure 1 provides an overview of the feature computation and its inputs.

**Figure 1:**
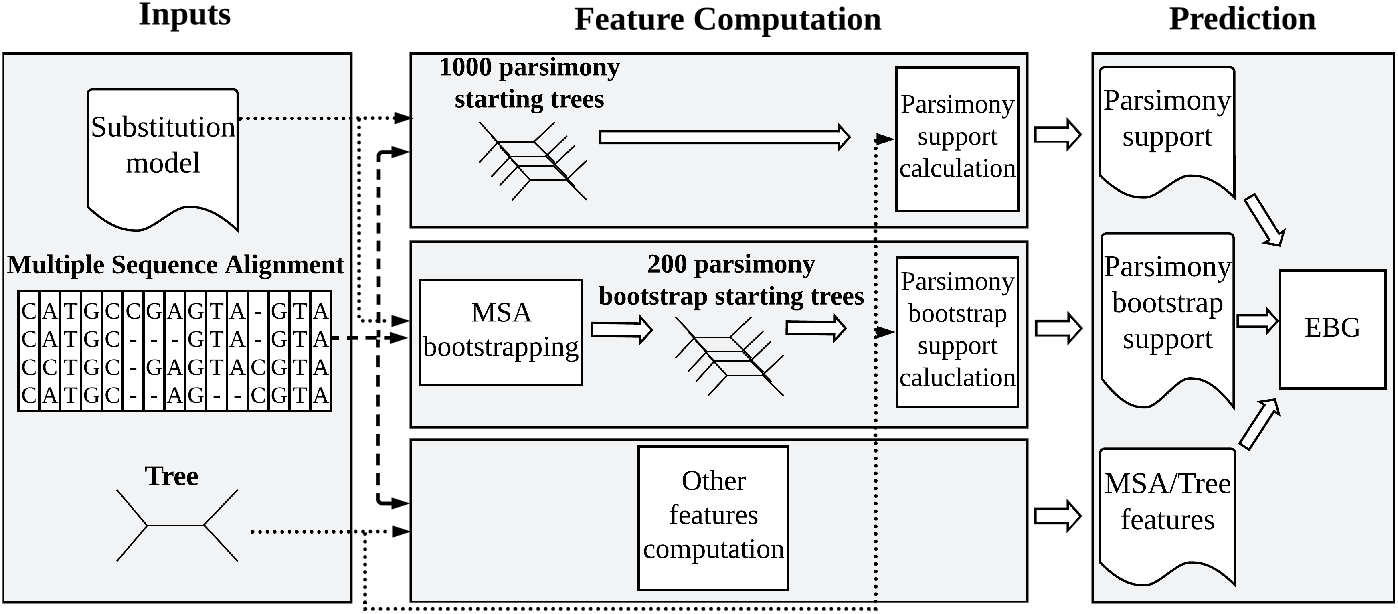
Overview of the EBG feature generation and prediction.

The PS is the frequency of occurrence of an inner branch in 1000 parsimony starting trees, we infer from the original MSA. According to exploratory experiments (see Supplementary Material Section 7), we set the number of inferred parsimony trees to 1000, as more than 1000 did not substantially improve prediction performance.

The PBS employs a procedure that is highly similar to the SBS and uses replicate MSAs. The only difference is, that PBS computes a parsimony starting tree for each bootstrap replicate, instead of inferring an ML tree, yielding the computation substantially faster. To capture the variance of the respective bootstrap tree space, we also use the mean normalized Robinson-Foulds (nRF) [26] distance between all PB trees as a feature. Again, based on exploratory experiments, inferring more than 200 PBs does not improve predictor performance (see Supplementary Material Section 7).

In theory, we could decrease the number of trees as a function of the Pythia difficulty score, since the tree space is less complex for lower Pythia difficulties. Hence, for lower Pythia difficulties, a smaller number of parsimony trees might be sufficient to approximate the SBS values. However, because this might introduce additional uncertainty in the prediction process, we decided to keep the number of parsimony trees for both the PBS and the PS features fixed.

We expected that the P(B)S values of branches adjacent to a focal inner branch of interest are indicative of its SBS value. Therefore, we also included summary statistics for the P(B)S values of respective child and parent branches as features. Additionally, we computed summary statistics over the per-site Parsimony Mutation count Scores (PMS) as another group of features. Finally, we use the mean closeness centrality [4] of the two subtrees connected to the focal inner branch. Closeness centrality quantifies how densely connected the tree’s nodes are to each other. By taking the closeness centrality ratio of those subtrees, we aimed to capture if the branch connects two subtrees of different densities. Another feature, we refer to as the branch length ratio bipartition, represents the ratio between the sums of branch lengths in the two subtrees induced by the focal branch.

The set of 23 features (see Table 1) we used for EBG is a subset of a larger set of over 150 features we experimented with. The Supplementary Material Section 2 comprises a detailed description of these features. We reduced this initial set of 150 features to 23 features via recursive feature elimination [11] and a scikit-learn random forest [24]. The aim was to determine a good trade-off between feature number and EBG performance for the following reasons. First, we strive to avoid unnecessary computations. Second, fewer features make the predictor more interpretable. Third, we aim to prevent overfitting our predictor to the training data. This can occur when training a predictor with too little training data and too many, unnecessary features [13]. In this case, the predictor will not generalize well to unseen inputs.

### 2.5 Predictor and Performance Metrics

According to preliminary experiments (cf. Supplementary Material Section 6), the best machine learning model choice for SBS prediction (both regression and classification) is the Light Gradient-Boosting Model (LightGBM) [16] tree-based boosting ensemble framework. To quantify the confidence of the EBG regression, we estimate the prediction uncertainty using quantile regression [17]. We use this approach to estimate the conditional quantiles of the SBS value. By training the model to predict the 5th and 10th quantile of the SBS value, we can provide lower bounds for the SBS value at 5% and 10%.

We optimized the hyperparameters of all models in 100 trials using the Optuna [2] hyperparameter optimization framework. Throughout the training process, we ensured that the TreeBASE MSAs we used were either exclusively contained in the training set or the test set. This guaranteed, that the inner branches of the respective ML tree are never split between the training and test set. Thereby we obtained an estimate of the predictors’ performance on entirely new, that is, unseen trees. We evaluated the performance of EBG using two distinct sets of metrics, one for the regression, and one for the classification step. In the following, we briefly describe both sets of metrics (see Supplementary Material Sections 3 and 4 for further details on their computation).

Botchkarev [5] proposes a regression metric typology, which we used to evaluate the EBG regressor. The typology describes regression performance metrics based on aggregation methods, distance measures, and normalization methods. Normalization is only necessary if we compare multiple predictions of different scales, which does not apply to EBG. We chose both, mean, and median as aggregation methods since the mean is sensitive to outliers, while the median is not. Additionally, we selected three distance measures. The squared distance between SBS values and the EBG prediction is sensitive to outliers. Furthermore, we selected the difference and the absolute difference between SBS and EBG values, which are not sensitive to outliers. This results in the following four metrics for evaluating the EBG regressor: the Mean Bias Error (MBE), the Mean Absolute Error (MAE), the Median Absolute Error (MdAE), and the Root Mean Squared Error (RMSE). We used the MBE to evaluate if the predictor is biased towards over- or underestimation. The MAE provides an outlier-sensitive, average performance of the predictor. The MdAE yields an outlier-insensitive performance metric. Finally, we used the RMSE as a measure that penalizes larger prediction errors more harshly.

The EBG classifier predicts class probabilities for the binary classification task, that is, the probability that an SBS value is below or above a specific SBS threshold. We set the decision boundary for the binary classification to 0.5. We evaluated the prediction accuracy of the EBG classifier using four metrics: the accuracy (Acc), the balanced accuracy (BAC), the F1-score (F1), and the Area Under the Curve (AUC) of the Receiver Operating Characteristic (ROC) curve. The Acc is easy to interpret and sensitive to class imbalances. The BAC is equivalent to the Acc with a weighing which is inversely proportional to the class prevalence. Therefore, the BAC is insensitive to class imbalances. The F1 is the harmonic mean between precision and recall, thus it penalizes false positives (FP) (erroneously predicting that the SBS value surpasses the SBS threshold) and false negatives (FN) as being equally unfavorable. The F1 is robust against class imbalances. In contrast to Acc and F1, the AUC allows for a classification performance evaluation without the need to set a class probability decision boundary, since it relies on the raw class probabilities.

## 3 Results

We evaluated the performance of the EBG regressor and classifier using different splits of our curated dataset of 1496 TreeBASE MSAs into training and testing MSAs. For both, the EBG regressor and classifier, we further tested the usage of EBG’s uncertainty measures for quantifying the reliability of its predictions. We also compared EBG with RB, UFBoot2, and SH-like aLRT regarding prediction quality, and we compared EBG against its fastest competitors UFBoot2 and SH-like aLRT in terms of time-to-completion and accumulated CPU time. Furthermore, we analyzed the importance of the prediction features for EBG.

### 3.1 EBG Regressor Performance Evaluation

The EBG regressor predicts three values: In addition to the central SBS point estimate, EBG provides two SBS predictions that correspond to two lower bounds. One lower bound has a 5%, the other a 10% predicted probability that the SBS value is below the respective bound. On a random subset of 232 MSAs, EBG’s central SBS estimate is highly correlated (mean Pearson correlation of *μ* = 0.91, *σ* = 0.05) with the SBS values. For a more detailed view, we refer to Supplementary Material Section 9. For the evaluation of the EBG regressor’s central SBS point estimate, we randomly sampled 20% of the 1496 MSAs as a holdout testing dataset and trained EBG on the remaining 80%. Table 2 summarizes the results of 10 such random holdouts. To assess the predictive power of EBG,

**Table 2:** EBG regression performance for 10 repeated random holdouts against a parsimony bootstrap support of 200 replicate trees as the baseline. To infer those replicate trees, we first obtain 200 replicate MSAs by sampling the original MSA site-wise with replacement. Subsequently, we infer the corresponding parsimony (starting) tree using RAxML-NG to calculate the 200 replicate trees.

| ***Metric*** | ***EBG*** ($\mu \pm \sigma$) | ***Baseline*** ($\mu \pm \sigma$) |
| --- | --- | --- |
| MBE | $0.6 \pm 0.3$ | $0.1 \pm 0.5$ |
| MAE | $8.3 \pm 0.2$ | $13.8 \pm 0.4$ |
| MdAE | $5.0 \pm 0.1$ | $8.6 \pm 0.5$ |
| RMSE | $12.8 \pm 0.2$ | $20.5 \pm 0.5$ |

we defined a baseline SBS value prediction based on the PBS of 200 replicates. If a branch is in the ML tree, but not in the PBS replicate tree set, we assigned a baseline prediction of zero to it, as this resembles the behavior of the SBS procedure. EBG outperformed this baseline across all four metrics. As the MBE indicates, the regressor exhibits no substantial systematic bias in either an over- or underestimation of the SBS values. The difference between MAE and MdAE, along with the higher RMSE value, suggests deviations of varying size between the EBG prediction and the true SBS value. The lower bound predictions of EBG can serve as a means to establish a bound for this prediction error.

Figure 2 illustrates, how the proximity of the lower bound predictions to the median prediction can be used to constrain the MdAE for the EBG median regression predictions. This approach effectively mitigates prediction uncertainty. As the lower bound predictions approach the median prediction, the MdAE decreases. Thus, the inspection of the distance between the lower bound and the median SBS predictions limits and quantifies the prediction uncertainty.

**Figure 2:**
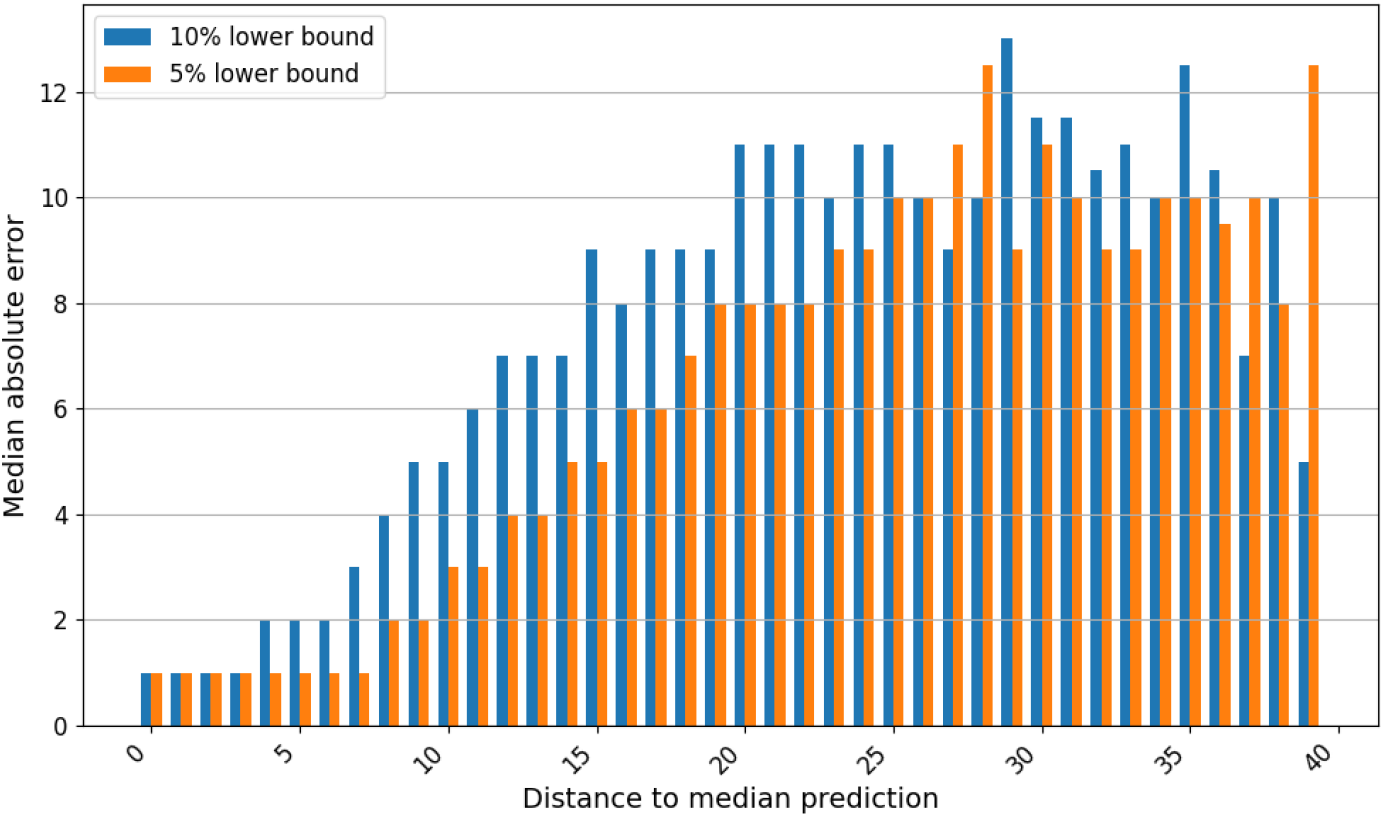
Relationship between the distance of the lower bound predictions to the median prediction and their influence on the MdAE.

### 3.2 EBG Classifier Performance Evaluation

In analogy to the EBG regressor, we evaluated the EBG classifier based on 10 repeated holdout sets of 20%. Table 3 summarizes the resulting performance metrics for varying SBS thresholds *t*. As baseline performance, we used the different SBS thresholds *t ∈ {*70, 75, 80, 85*}* on the PBS of 200 replicates. The baseline again is zero for all branches, that are present in the ML tree, but not in the PBS replicate trees. EBG outperformed the baseline for every metric.

**Table 3:**
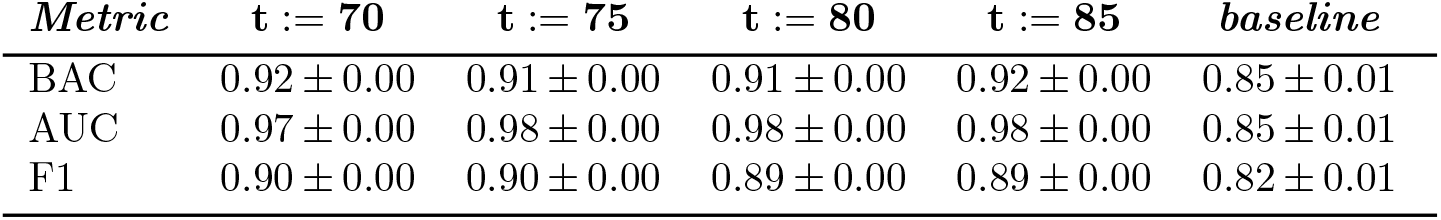
EBG classifier performance for different decision boundaries *t*. We indicate the mean and standard deviation of 10 repeated random holdouts against a baseline consisting of the PBS of 200 replicates.

| ***Metric*** | ***t := 70*** | ***t := 75*** | ***t := 80*** | ***t := 85*** | ***baseline*** |
| --- | --- | --- | --- | --- | --- |
| BAC | $0.92 \pm 0.00$ | $0.91 \pm 0.00$ | $0.91 \pm 0.00$ | $0.92 \pm 0.00$ | $0.85 \pm 0.01$ |
| AUC | $0.97 \pm 0.00$ | $0.98 \pm 0.00$ | $0.98 \pm 0.00$ | $0.98 \pm 0.00$ | $0.85 \pm 0.01$ |
| F1 | $0.90 \pm 0.00$ | $0.90 \pm 0.00$ | $0.89 \pm 0.00$ | $0.89 \pm 0.00$ | $0.82 \pm 0.01$ |

In analogy to the lower bound prediction for the EBG regressor, EBG also provides a prediction uncertainty measure for its classifier. As the EBG classifier solves a binary classification problem, with one class defined as *SBS > t* and the other as *SBS ≤ t*, we can leverage the Shannon entropy [27] of the two class probabilities to obtain a prediction uncertainty measure *u ∈* [0, 1]. Here, *u* = 0 represents absolute certainty, and *u* = 1 corresponds to absolute uncertainty. Figure 3 provides an overview of the relationship between EBG classifiers with different SBS thresholds *t* and their Acc with increasing prediction uncertainty. Especially for the smaller uncertainties, a high class imbalance implies a better classification within the uncertainty interval of concern. Therefore, we decided not to compensate for class imbalances by using the BAC and used the Acc for this analysis instead. For cases with low *u ∈* [0.1, 0.3], the prediction Acc consistently remains at or above 90% across all SBS thresholds *t*. For moderate *u ∈* [0.4, 0.6], the Acc typically falls within the 80% to 90% range. We only observe prediction accuracies below 70% for *u >* 0.8.

**Figure 3:**
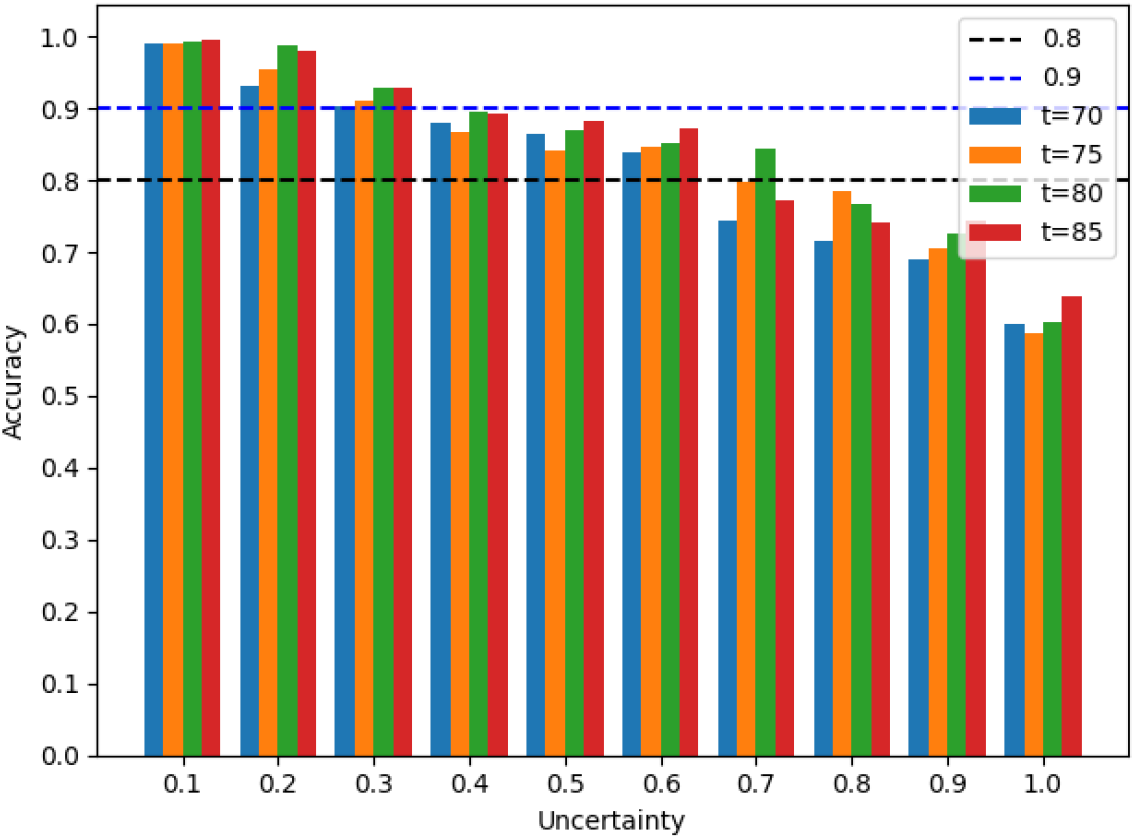
Relationship between the prediction Acc of EBG classifiers and their prediction uncertainty for varying SBS thresholds *t*.

In practical terms, when using EBG, we have the flexibility to determine an acceptable level of uncertainty to attain confident predictions. Thereby, users can tailor the approach to their specific needs and preferences.

To demonstrate the performance of EBG on an edge case, we predicted the SBS values for an MSA that was shown to be difficult to analyze in terms of phylogenetic inference. Morel et al. [22] demonstrate the difficulties of obtaining a reliable phylogeny using ML inference on a set of SARS-CoV-2 genome sequences. This difficulty is caused by the combination of a large number of sequences and a relatively low mutation rate, resulting in numerous branches with low bootstrap values. For our experiment, we used a set of 1654 complete SARS-CoV-2 genomes, each with a length of 29 800 base pairs. We employed 100 RAxML-NG searches to determine the ML tree with the highest log-likelihood. We then performed an SBS run with 1000 replicates to establish the ground truth SBS values. This analysis shows, that only 13.9% of the inner branches of the ML tree yield an SBS value greater than 70 indicating an overall low support of the ML tree.

Predicting the branch support using our EBG regressor results in an overall good performance, with an MAE of 3, an MdAE of 0, and an RMSE of 9. Meanwhile, the EBG classifier, with an SBS threshold of *t* := 70, achieved a BAC of 0.82, an F1 of 0.77, and an AUC of 0.82. Even on an MSA that is known to be difficult-to-analyze, EBG can provide a good estimate of the SBS values for the corresponding ML tree. The prediction of the bootstrap support, using a mid-class laptop equipped with 4 cores and 8 GB of memory, has a time-to-completion of approximately three hours. In contrast, computing the ground truth SBS values took 35 hours, utilizing 10 nodes of a large computing cluster, each equipped with an Intel Xeon Gold 6230 (20 cores, 2.1 GHz) and 96 GB memory. On the same amount of computing nodes the MPI-based UFBoot2 analysis only took 31 minutes.

Another dataset that is notoriously difficult to analyze is the internal transcribed spacer (ITS) 354 [9]. ITS 354 is a short alignment (348 MSA sites) extracted from the ITS genes from 354 maple tree genomes. Predicting the SBS values using the EBG regressor results in an MAE of 6, an MdAE of 4, and an RMSE of 9. Hence, on this difficult MSA EBG also yields a good performance.

### 3.3 Performance Comparison with UFBoot2, SH-like aLRT, and Rapid Bootstrap

EBG, UFBoot2, and SH-like aLRT branch support values have all different interpretations. EBG approximates the SBS which is conservative, whereas UFBS values are unbiased [21]. SH-like aLRT is also conservative but not in the same way as SBS, reasonable thresholds for SH-like aLRT can be between 0.8 and 0.9 [10]. Therefore, to compare all three branch supports against each other, we needed a reliable ground truth phylogeny. Consequently, we performed the following performance comparison using simulated MSAs. We simulated a total of 979 DNA MSAs without gaps (i.e., without simulating indel events) based on TreeBASE trees using AliSim [19] under the GTR+G model. The corresponding tree (true tree) of the simulated MSAs served as the ground truth for our experiment. We compared the fraction of branches in the true tree for each branch support value of EBG, UFBoot2, and SH-like aLRT. A similar approach was used for the evaluation of the original UFBoot by Minh et al. [21]. For the following analyses, we used the IQ-TREE2 [21] implementation of the SH-like aLRT. The commands we used are part of Supplementary Material Section 1. Figure 4 summarizes the results of our comparison analyses. An ideal branch support measure would yield the unbiased probability of the branch being in the true tree (dashed, red line). In our experiments, all three tools are too liberal with their estimation of the true branch probability. As already observed by Minh et al. [21], low SH-like aLRT values (*<* 50) are not informative concerning the true probabilities. For larger values, the SH-like aLRT behaves similarly to EBG. EBG is fairly unbiased for low support values (*<* 60). For branch supports *>* 60 it tends to overestimate the true probability of the branches being present in the true tree. In this experiment, UFBoot2 overestimated the true branch probability the most. Besides all three tools being overconfident in predicting the true support, EBG is the closest to the ideal branch support value line.

**Figure 4:**
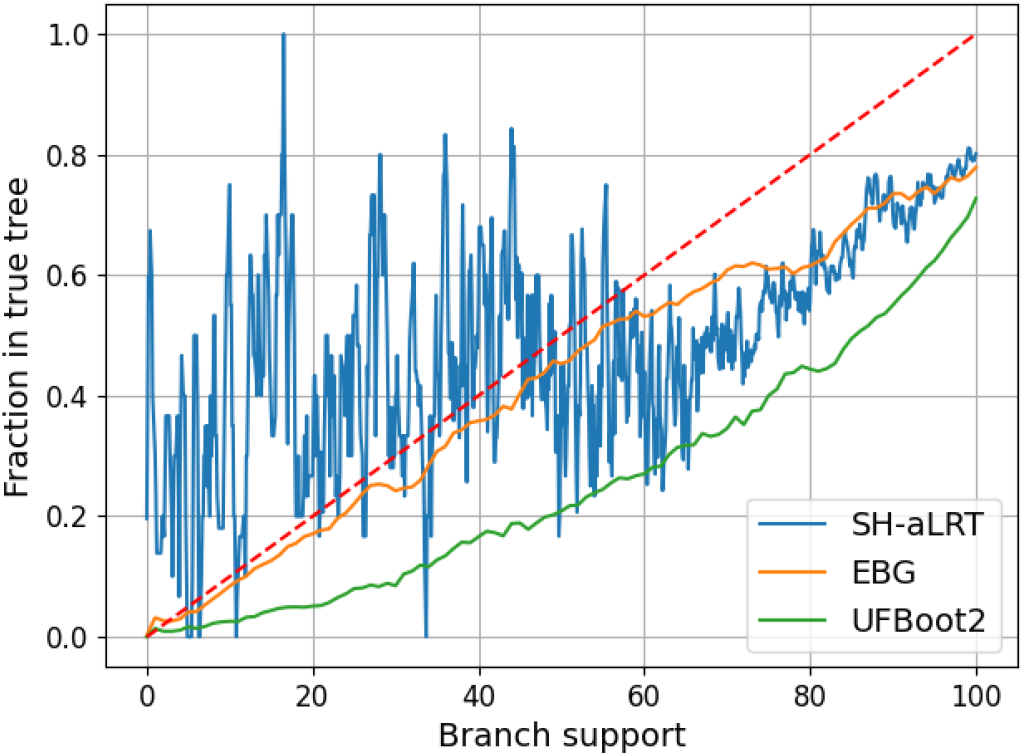
Moving average with window size five of the fraction of branches in the true tree for all branch support values.

We also compared the performance of the different tools as a function of the Pythia difficulty of the simulated MSAs. We computed the BAC for specific branch support thresholds and compared them across all tools. The results suggest that EBG is best able to deal with varying MSA difficulties. We provide the detailed analyses in Supplementary Material Section 5.

Finally, we compared EBG directly with RB on empirical MSAs. This is possible since RB is highly correlated with the SBS values [29]. We randomly selected 220 MSAs (20% AA, 80% DNA) from TreeBASE and computed the ground truth SBS for each branch based on 1000 bootstrap replicates using RAxML-NG. Table 4 summarizes the respective classification and regression metrics for EBG and RB.

**Table 4:** Performance evaluation of the EBG regressor, EBG classifier with SBS threshold *t* := 0.80, and RB.

| *Tool* | *MBE* | *MAE* | *MdAE* | *RMSE* | *BAC* | *F1* | *AUC* |
| --- | --- | --- | --- | --- | --- | --- | --- |
| EBG | 0.0 | 8.7 | 6.0 | 12.8 | 0.89 | 0.87 | 0.89 |
| EBG* | 0.1 | 7.1 | 4.0 | 11.1 | 0.95 | 0.94 | 0.95 |
| RB | 0.0 | 4.5 | 2.0 | 8.0 | 0.97 | 0.96 | 0.97 |

\* EBG with consideration of prediction uncertainty

Based on our experimental findings, RB achieves the highest Acc when predicting SBS values. EBG’s prediction performance falls short compared to RB, as evidenced by a higher MdAE (RB: 2.0, EBG: 6.0) and a lower BAC (RB: 0.96, EBG: 0.89). However, if we also use EBG’s prediction uncertainty measures, the performance becomes comparable, in particular for the EBG classifier.

Note that EBG* in Table 4 summarizes the results for an uncertainty filtering of the predictions we conducted for both, the regression, and classification tasks. In the regression scenario, our focus is on branches where we expect the MdAE to be less than or equal to 8 (as Figure 2 depicts), that is, a 5% lower bound distance of 23 or less. This uncertainty filtering results in the exclusion of 28% of predictions that we deem too uncertain for consideration. For classification, we restrict our attention to predictions with an uncertainty level *u ≤* 0.7 (as Figure 3 depicts). This leads to excluding 21% of the predictions, that we consider as being too uncertain. While this approach may not provide predictions for every branch, it effectively constrains prediction errors. It represents a trade-off between the number of predictions that we consider to be trustworthy and the level of certainty of those predictions.

In addition to the above accuracy analyses, we conducted a comparison of the time-to-completion of EBG with UFBoot2 and the SH-like aLRT (using IQ-TREE2) as fastest competitors using empirical datasets. The SBS computation with RAxML-NG, UFBoot2 as well as the a-LRT implementation of IQ-TREE2 can use multiple threads. Since the independent parsimony tree inferences necessary for the feature computation of EBG can also be parallelized straightforwardly, we performed our benchmark on a reference machine using multiple threads. This reference machine is equipped with an Intel Xeon Platinum 8260 Processor (48 physical cores, 2.4 GHz) and 754 GB memory. RAxML-NG and IQ-TREE2 provide the option to automatically determine the optimal number of threads for a given MSA, and we used this feature in both tools with up to a total of 60 threads. Figure 5 summarizes the results of the benchmark on the 220 empirical TreeBASE MSAs we used for the EBG/RB comparison. For the sake of simplicity, we define MSA size as the product of the number of sequences and the number of site patterns (unique MSA sites). Furthermore, we separately depict run times for AA (dashed lines) and DNA datasets (straight lines) to assess potential differences between data types. Since EBG requires an existing phylogenetic tree and substitution model parameters as input, we included the inference time of adaptive RAxML-NG [31] in the time-to-completion of the EBG prediction (EBG + inference). We observed that over all 220 MSAs, EBG accounts for 19% and the ML inference for 81% of the total time-to-completion. SH-like aLRT and UFBoot2 exhibit a similar time-to-completion, both for DNA, and AA MSAs. For 97% of the datasets (DNA and AA), EBG outperformed UFBoot2 in terms of time-to-completion, with an average speedup of 9.4 (*σ* = 5.5). Considering MSAs of size *≥* 200 000, EBG yielded an average speedup of 2.8 in comparison to UFBoot2. With increasing MSA sizes, the time-to-completion differences between UFBoot2 and EBG gradually decrease for AA datasets. We do not observe an analogous trend for DNA data.

**Figure 5:**
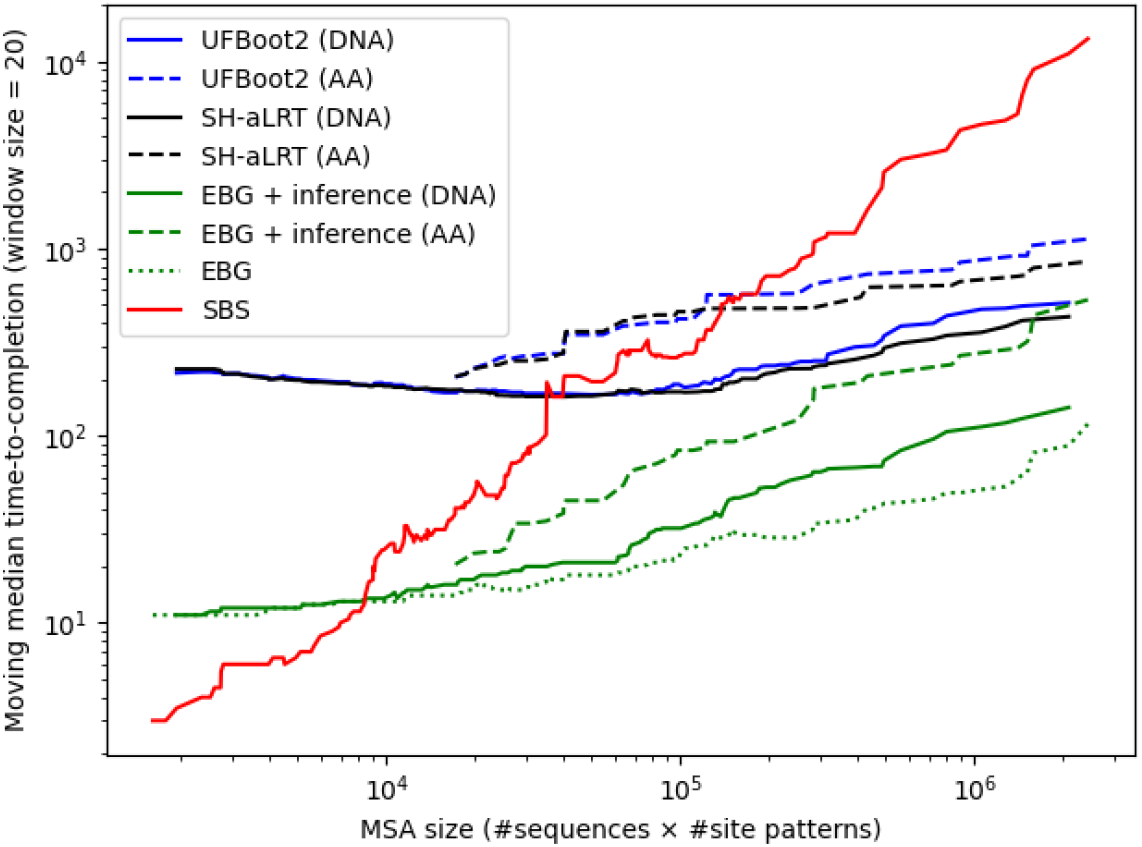
Time-to-completion comparison for datasets of varying size with a moving average of window size 20.

Additionally, we used a random sample of 84 MSAs from the total of 220 MSAs to assess the disparity in accumulated CPU time between EBG and UFBoot2. In the median, running EBG along with an adaptive RAxML-NG ML tree search requires 28% less accumulated CPU time in comparison to the corresponding UFBoot2 execution. However, summed over all 84 MSAs EBG including the inference take 57% more accumulated CPU time (65 137 vs. 43 090 seconds) than UFBoot2. This suggests that there are some outliers where UFBoot2’s time-to-completion is substantially smaller than EBG’s time-to-completion when we include the ML inference time in the calculation. According to our analysis, this seems to be primarily the case with MSAs having a Pythia difficulty of *≥* 0.5 (see Supplementary Material Section 8 for a detailed visualization). Furthermore, we observe, that on the same 84 MSAs EBG only accounts for 4% of the accumulated CPU time while the remaining 96% are required for the adaptive RAxML-NG tree inference that is required as an input for EBG.

We conclude that it is challenging to devise a fair time-to-completion comparison between EBG and UFBoot2 as for EBG, we are undecided if the ML inference time should be included or not. In contrast, for UFBoot2 it represents an intrinsic requirement for the computation of support values as the ML search and the support calculations are necessarily intertwined. Hence, the above time comparisons reflect, to a large extent, a time comparison between the adaptive RAxML-NG and IQ-Tree ML search algorithms.

### 3.4 Feature Importances

Table 5 lists the five most important features of the EBG regressor. The feature importance quantifies to which extent a feature contributes to achieving an improved prediction during training. According to the feature importances of EBG, the PBS feature with a feature importance of 82.2% is by far the most important one for predicting SBS values. We interpret the substantial importance of the PBS features with an analogy to ensemble methods in machine learning. These methods aggregate numerous weak learners, to create a robust, strong one. Similarly, we use the contributions of multiple “weak” parsimony inferences, to obtain a precise estimate of the SBS values. See Supplementary Material Section 2 for a more detailed overview of all prediction features and their importance.

**Table 5:** Overview of the five most important features that EBG uses for the prediction and their respective importance in percent.

| *Feature* | *Importance in %* |
| --- | --- |
| PBS | 82.2 |
| PS | 3.1 |
| Normalized branch length | 2.0 |
| # child inner branches | 1.7 |
| Skewness PBS | 1.5 |

## 4 Discussion and Conclusion

In this paper, we introduced EBG, a novel machine learning-based approach for predicting SBS values of phylogenies.

We observe that RB remains the most accurate approximation method for the direct prediction of SBS values. We demonstrate how the EBG uncertainty measures can help to reduce the accuracy gap to RB. Filtering the predictions of the EBG classifier to an uncertainty level *u ≤* 0.7 closes this accuracy gap based on our experiments. EBG including the ML tree inference time requires substantially lower time-to-completion compared to the major competitor UFBoot2 with an average speedup of 9.4 (*σ* = 5.5). This speedup comes at the cost of an increase in total accumulated CPU time summed over all test MSAs of 57% due to outliers.

On 979 simulated MSAs, EBG, UFBoot2, and SH-like aLRT are generally too liberal and provide mixed results. Besides that, EBG yields support predictions that are closest to the ideal branch support value.

Currently, EBG implements the sampling part of the PB procedure in Python. The PB is a performance bottleneck that substantially contributes to the prediction time and often accounts for up to 70% of the overall time-to-completion of EBG excluding the ML inference time. Furthermore, EBG performs 204 RAxML-NG calls: One for each of the 200 parsimony bootstrap inferences, one for the parsimony inference, two for the support computation of both, and one for the nRF distance computation. Additionally, we store and retrieve the results of those calls in individual files, posing an I/O overhead. Unifying EBG’s feature computation and the prediction as a command implemented within the RAxML-NG tool would likely result in a substantial speedup and streamline the entire prediction process. Since RAxML-NG is developed in our lab, the integration of EBG into RAxML-NG constitutes future work.

While EBG successfully establishes lower bounds for the EBG regression, we were not able to devise a prediction for the upper bound of SBS values. In our experiments, the attempts to optimize for any upper bound were unsuccessful, since they converged to the trivial upper bound of an SBS of 100. An upper bound for EBG regression would be useful to construct a comprehensive prediction interval. Future research could focus on the development of a method or model that is capable of reliably estimating upper bounds. A more informative prediction interval will be highly beneficial for assessing the range of potential SBS values.

Finally, we note that parsimony-based methods may experience a renaissance, since as we show here and as already demonstrated by the Pythia tool, they constitute the by far most important and computationally inexpensive feature for conducting predictions about ML method results.

## Acknowledgements

This work was financially supported by the Klaus Tschira Foundation and by the European Union (EU) under Grant Agreement No 101087081 (Comp-Biodiv-GR).

