## Supplementary Material for "Predicting Phylogenetic Bootstrap Values via Machine Learning"

March 2024

### 1 Command References

| ***Tool*** | ***Version*** | ***Commands*** |
|---|---|---|
| RAxML | 8.2.12 | **Rapid Bootstrap**: raxmlHPC-PTHREADS -T 60 -m GTRGAMMA/PROTGAMMALG -s msa_filepath -# 1000 -p 12345 -x 12345<br>**Rapid Bootstrap (MPI)**: mpirun raxmlHPC-MPI -m GTRGAMMA/PROTGAMMALG -s msa_filepath -# 10 -p 12345 -x 12345 |
| RAxML-NG | 1.1.0 | **Search**: raxml-ng −−adaptive −−msa msa_filepath −−model GTR+G/LG+G −−threads auto{60}<br>**SBS**: raxml-ng −−bootstrap −−model model_filepath −−bs−trees 1000 −−msa msa_filepath<br>**SBS (MPI)**: mpirun raxml-ng −−bootstrap −−model model_filepath −−bs−trees 1000 −−msa msa_filepath −−workers 50 |
| IQ-TREE2 | 2.2.2.7 | **UFBoot2**: iqtree2 -m GTR+G/LG+G -s msa_filepath -B 1000 -T AUTO –threads-max 60<br>**UFBoot2 (MPI)**: mpirun iqtree2-mpi -m GTR+G/LG+G -s msa_filepath -B 1000 -T AUTO –threads-max 60<br>**SH-like aLRT**: iqtree2 -m GTR+G/LG+G -s msa_filepath -alrt 1000 -T AUTO –threads-max 60 |

Table 1: Used commands for each tool

### 2 Features

| *Feature* | *Importance (%)* |
|---|---|
| Parsimony bootstrap support (PBS) | 82.2 |
| Parsimony support (PS) | 3.1 |
| Normalized branch length | 2.0 |
| # child inner branches | 1.7 |
| Skewness PBS | 1.5 |
| Mean Robinson-Foulds distance PB | 1.1 |
| Mean parsimony substitution frequency (PSF) | 1.1 |
| Branch length | 1.0 |
| Max. PSF | 1.0 |
| Coefficient of variation PSF | 0.7 |
| Max. PBS children* | 0.6 |
| Mean PBS parents | 0.6 |
| Max. PS children* | 0.5 |
| Branch number (ordered by level-order traverse) | 0.5 |
| Skewness PSF | 0.5 |
| Branch length ratio bipartition | 0.5 |
| Max. PS children* | 0.3 |
| Min. PS children | 0.3 |
| Std. PBS parent branches | 0.3 |
| Std. PBS child branches | 0.3 |
| Mean closeness centrality bipartition ratio | 0.3 |
| Min. PS children* | 0.2 |
| Min. PBS children* | 0.1 |

*: weighted by branch length

Table 2: Overview of the subset of features used for the prediction and their final feature importance for the regressor. We obtained this subset via Recursive Feature Elimination.

- **Parsimony Support (PS)**
  We compute the PS using the −−start-option of RAxML-NG for generating 1000 parsimony starting trees. We calculate the support using RAxML-NG as well. More than 1000 parsimony starting trees do not yield a better predictor performance according to our experiments.

- **Parsimony Bootstrap Support (PBS)**
  We generate a PB by resampling the columns of the MSA with replacement. For each of the sampled MSAs, we compute the corresponding parsimony starting tree using the −−start-option in RAxML-NG. We infer a total of 200 PB trees. More than 200 PB trees do not yield a better predictor performance according to our experiments.

- **Normalized branch length**
  We normalize the branch lengths by the total sum of branch lengths of the tree.

- **# child inner branches**
  The number of inner branches in the subtree below the branch.

- **Skewness PBS**
  The skewness of the PBS of all inner branches of the input tree.

- **Mean Robinson-Foulds distance PB**
  The mean Robinson-Foulds distance of the trees generated by the PB procedure.

- **Parsimony substitution frequency (PSF)** [**statistic**]
  We compute the number of parsimony substitutions per site using the tree and the MSA. Afterwards, we calculate the summary statistics.

- [**Min**|**Max**|**Mean**|**Std**|**Skewness**] **P(B)S** [**children**|**parents**][*****]
  We compute the statistics over the P(B)S for the inner branches below (children) or above (parents) the branch. As indicated by *, we also weigh the P(B)S values of the child/parent inner branches by their branch length for some features.

- **Mean closeness centrality bipartition ratio**
  We split the tree into two parts at the branch. We then transform the trees into graphs with networkx [2] and calculate their mean closeness centrality. We define the closeness centrality of node $i$ as in Equation (1) [1] with $N$ being the number of nodes in a graph and $d(x,y)$ as the shortest branch distance between two nodes.

  $$C(i) = \frac{1}{\sum_{i=1}^{N} d(x,i)} \tag{1}$$

  By dividing the smaller by the larger closeness centrality we obtain the final feature.

### 3 Regression Metrics

#### 3.1 Mean Bias Error

$$MBE = \frac{1}{N}\sum_{i=1}^{N}(y_i - \hat{y_i}) \tag{2}$$

##### 3.1.1 Mean/Median Absolute Error

$$MAE = \frac{1}{N}\sum_{i=1}^{N}(|y_i - \hat{y_i}|) \tag{3}$$

##### 3.2 (Root) Mean Squared Error

$$MSE = \frac{1}{N}\sum_{i=1}^{N}(y_i - \hat{y_i})^2 \tag{4}$$

$$RMSE = \sqrt{MSE} = \sqrt{\frac{1}{N}\sum_{i=1}^{N}((y_i - \hat{y_i})^2)} \tag{5}$$

#### 4 Classification Metrics

##### 4.1 Accuracy

$$Acc = \frac{TP + FP}{TP + FP + TN + FN} \tag{6}$$

where $TP$ are true and $FP$ false positives (SBS threshold $t$ exceeded). We define $FP$ and $FN$ analogously.

##### 4.2 Balanced Accuracy

$$BAC = \frac{TP/P + TN/N}{2} \tag{7}$$

where $P$ are positives and $N$ are negatives.

##### 4.3 F1-score

$$F1 = \frac{2 \times \text{precision} \times \text{recall}}{\text{precision} + \text{recall}} \tag{8}$$

$$\text{precision} = \frac{\text{TP}}{\text{FP} + \text{TP}} \tag{9}$$

$$\text{recall} = \frac{\text{TP}}{\text{FN} + \text{TP}} \tag{10}$$

##### 4.4 ROC-AUC-score

It quantifies the area under the ROC (receiver operating characteristic) curve (AUC). In contrast to Acc and the F1, it allows for evaluation without the need to set a class probability decision boundary, since it relies on the raw class probabilities. The best binary classifier yields an AUC of 1, while a random classifier has an AUC of 0.5.

#### 5 Tool Balanced Accuracy and MSA Difficulty

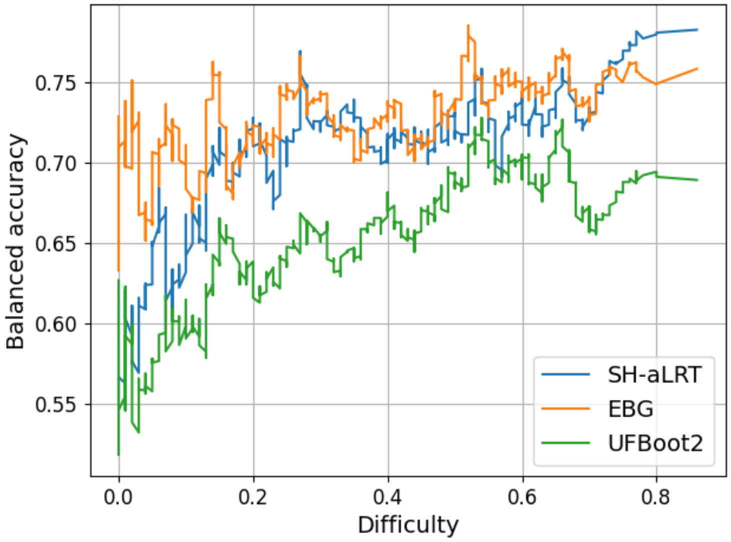


Figure 1: Balanced accuracy comparison of EBG, UFBoot2, and SH-like aLRT concerning Pythia difficulty.

We compared the BAC of the three tools in predicting whether a branch is in the true tree or not. We set the decision threshold for EBG and SH-like aLRT to 80, as they behave similarly on the simulated MSAs. For UFBoot2 we set the threshold for predicting the branch is in the true tree to 95. According to the authors, this yields a 5% false positive bound [3]. It is difficult to compare the BAC values of the three tools due to the definition of individual thresholds. At least with this set of thresholds, EBG yields the most consistent performance on the simulated MSAs.

#### 6 Model Comparison

| ***Tool*** | ***MAE*** | ***BAC*** |
|---|---|---|
| Logistic/Ridge Regression | $10 \pm 0.2$ | $0.89 \pm 0.00$ |
| Random Forest | $12 \pm 1.6$ | $0.89 \pm 0.00$ |
| LightGBM | $8.3 \pm 0.2$ | $0.91 \pm 0.00$ |

Table 3: Performance of different machine learning models on the regression and classification task formulation. The table shows the mean and standard deviation of the metrics based on 10 repeated random holdouts of size 20%.

### 7 Number of Parsimony Trees

| ***Configuration*** | ***MdAE*** | ***RMSE*** |
|---|---|---|
| 100 parsimony bootstrap starting trees | 7.9 | 13.8 |
| 200 parsimony bootstrap starting trees | 7.6 | 13.1 |
| 500 parsimony bootstrap starting trees | 7.5 | 13.0 |
| 100 parsimony starting trees | 9.3 | 14.9 |
| 1000 parsimony starting trees | 8.5 | 14.2 |
| 10000 parsimony starting trees | 8.4 | 14.0 |

Table 4: Performance comparison of the EBG regressor with different numbers of parsimony (bootstrap) trees as the basis for the P(B)S features. The table shows the mean of the metrics based on 10 repeated random holdouts of size 20%. As 500 PB trees did not yield a significantly better MdAE compared to 200 PB trees, we chose to use 200 PB trees for a shorter time-to-completion. The same rationale led to the choice of 1000 parsimony starting trees.

### 8 Accumulated CPU Time Comparison

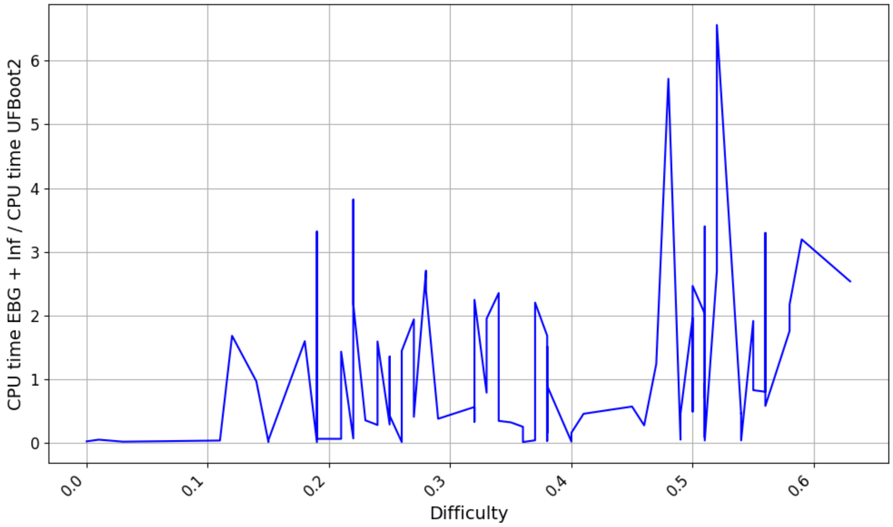


Figure 2: Ratio of the EBG and inference accumulated CPU time against the UFBoot2 accumulated CPU time on 84 MSAs.

### 9 Pearson Correlation EBG Regressor

| MSA | Pearson correlation | p-value |
|---|---|---|
| 10098_1 | 0.91 | 0.0 |
| 10115_3 | 0.88 | 0.0 |
| 10118_0 | 0.94 | 0.0 |
| 10169_0 | 0.95 | 0.0 |
| 10169_2 | 0.93 | 0.0 |
| 10196_0 | 0.94 | 0.0 |
| 10264_0 | 0.9 | 0.0 |
| 10268_3 | 0.96 | 0.0 |
| 10270_3 | 0.83 | 0.0 |
| 10271_5 | 0.92 | 0.0 |
| 10436_0 | 0.94 | 0.0 |
| 10436_8 | 0.84 | 0.0 |
| 10454_1 | 0.94 | 0.0 |
| 10542_1 | 0.93 | 0.0 |

| | | |
|---|---|---|
| 10562_0 | 0.94 | 0.0 |
| 10568_0 | 0.95 | 0.0 |
| 10629_0 | 0.96 | 0.0 |
| 10652_0 | 0.92 | 0.0 |
| 10703_5 | 0.95 | 0.0 |
| 10714_0 | 0.95 | 0.0 |
| 10727_0 | 0.92 | 0.0 |
| 10749_0 | 0.97 | 0.0 |
| 10782_0 | 0.95 | 0.0 |
| 10791_9 | 0.87 | 0.0 |
| 10801_0 | 0.67 | 0.0 |
| 10856_1 | 0.87 | 0.0 |
| 10949_2 | 0.83 | 0.0 |
| 10983_0 | 0.78 | 0.0 |
| 10986_0 | 0.91 | 0.0 |
| 11032_4 | 0.9 | 0.0 |
| 11331_0 | 0.89 | 0.0 |
| 11376_0 | 0.94 | 0.0 |
| 11487_7 | 0.95 | 0.0 |
| 11712_1 | 0.92 | 0.0 |
| 11768_1 | 0.93 | 0.0 |
| 11777_0 | 0.86 | 0.0 |
| 11783_0 | 0.93 | 0.0 |
| 11966_0 | 0.83 | 0.0 |
| 11972_1 | 0.94 | 0.0 |
| 11988_0 | 0.88 | 0.0 |
| 11991_4 | 0.95 | 0.0 |
| 12013_0 | 0.9 | 0.0 |
| 12020_0 | 0.95 | 0.0 |
| 12165_2 | 0.92 | 0.0 |
| 12306_1 | 0.91 | 0.0 |
| 12306_13 | 0.97 | 0.0 |
| 12334_0 | 0.93 | 0.0 |
| 12339_0 | 0.93 | 0.0 |
| 12493_1 | 0.91 | 0.0 |
| 12493_7 | 0.86 | 0.0 |
| 12717_2 | 0.93 | 0.0 |
| 12746_0 | 0.91 | 0.0 |
| 12855_2 | 0.95 | 0.0 |
| 13184_0 | 0.89 | 0.0 |
| 13664_0 | 0.9 | 0.0 |
| 13801_2 | 0.92 | 0.0 |
| 13808_7 | 0.93 | 0.0 |

| | | |
|---|---|---|
| 13815_2 | 0.97 | 0.0 |
| 13887_0 | 0.86 | 0.0 |
| 13909_0 | 0.93 | 0.0 |
| 13909_1 | 0.93 | 0.0 |
| 13985_6 | 0.92 | 0.0 |
| 14035_0 | 0.94 | 0.0 |
| 14151_0 | 0.98 | 0.002 |
| 14188_0 | 0.91 | 0.0 |
| 14232_0 | 0.83 | 0.0 |
| 14244_0 | 0.7 | 0.0 |
| 14504_0 | 0.94 | 0.0 |
| 14526_0 | 0.94 | 0.0 |
| 14534_25 | 0.95 | 0.0 |
| 14643_2 | 0.92 | 0.0 |
| 14663_0 | 0.85 | 0.0 |
| 14688_0 | 0.85 | 0.033 |
| 14688_13 | 0.92 | 0.0 |
| 14688_23 | 0.92 | 0.0 |
| 14725_5 | 0.95 | 0.0 |
| 14725_7 | 0.96 | 0.0 |
| 14833_4 | 0.94 | 0.0 |
| 14923_0 | 0.95 | 0.0 |
| 14954_0 | 0.9 | 0.0 |
| 14959_2 | 0.96 | 0.0 |
| 15019_5 | 0.89 | 0.0 |
| 15021_3 | 0.92 | 0.0 |
| 15021_7 | 0.94 | 0.0 |
| 15039_2 | 0.87 | 0.0 |
| 15097_1 | 0.93 | 0.0 |
| 15179_2 | 0.95 | 0.0 |
| 15253_0 | 0.82 | 0.0 |
| 15306_5 | 0.93 | 0.0 |
| 15368_0 | 0.91 | 0.0 |
| 15427_0 | 0.91 | 0.0 |
| 15635_1 | 0.89 | 0.0 |
| 15636_0 | 0.82 | 0.0 |
| 15639_0 | 0.87 | 0.0 |
| 15669_9 | 0.92 | 0.0 |
| 15682_0 | 0.92 | 0.0 |
| 15769_0 | 0.94 | 0.0 |
| 15828_0 | 0.95 | 0.0 |
| 15861_2 | 0.93 | 0.0 |
| 15908_1 | 0.86 | 0.0 |

| | | |
|---|---|---|
| 16007_0 | 0.87 | 0.0 |
| 16009_1 | 0.89 | 0.0 |
| 16105_0 | 0.88 | 0.0 |
| 16141_1 | 0.78 | 0.041 |
| 16190_2 | 0.94 | 0.0 |
| 16269_0 | 0.92 | 0.0 |
| 16313_11 | 0.94 | 0.0 |
| 16453_0 | 1.0 | 1.0 |
| 16629_0 | 0.88 | 0.0 |
| 16632_2 | 0.96 | 0.0 |
| 16637_2 | 0.97 | 0.0 |
| 16675_0 | 0.88 | 0.0 |
| 16737_0 | 0.84 | 0.0 |
| 16748_0 | 0.84 | 0.0 |
| 16785_1 | 0.9 | 0.0 |
| 16855_2 | 0.97 | 0.0 |
| 17014_0 | 0.9 | 0.0 |
| 17168_0 | 0.95 | 0.0 |
| 17390_1 | 0.92 | 0.0 |
| 17443_0 | 0.92 | 0.0 |
| 17594_11 | 0.93 | 0.0 |
| 17594_13 | 0.95 | 0.0 |
| 17666_0 | 0.89 | 0.0 |
| 17723_0 | 0.94 | 0.0 |
| 17749_1 | 0.9 | 0.0 |
| 17761_0 | 0.91 | 0.0 |
| 17774_4 | 0.97 | 0.0 |
| 17791_0 | 0.85 | 0.0 |
| 17814_0 | 0.93 | 0.0 |
| 17878_9 | 0.85 | 0.0 |
| 17885_1 | 0.96 | 0.0 |
| 17896_31 | 0.92 | 0.0 |
| 18077_0 | 0.94 | 0.0 |
| 18131_0 | 0.84 | 0.0 |
| 18218_1 | 0.96 | 0.0 |
| 18258_2 | 0.92 | 0.0 |
| 18438_0 | 0.75 | 0.003 |
| 18448_0 | 0.89 | 0.0 |
| 18465_0 | 0.94 | 0.0 |
| 18638_0 | 0.87 | 0.0 |
| 18638_1 | 0.92 | 0.0 |
| 18654_0 | 0.95 | 0.0 |
| 18850_2 | 0.62 | 0.574 |

| | | |
|---|---|---|
| 18883_0 | 0.92 | 0.0 |
| 19060_0 | 0.96 | 0.0 |
| 19447_0 | 0.91 | 0.0 |
| 19466_3 | 0.92 | 0.0 |
| 19509_1 | 0.99 | 0.0 |
| 19579_0 | 0.91 | 0.0 |
| 19740_5 | 0.82 | 0.0 |
| 19782_3 | 0.78 | 0.0 |
| 19797_0 | 0.95 | 0.0 |
| 19889_1 | 0.89 | 0.0 |
| 19925_0 | 0.96 | 0.0 |
| 19925_6 | 0.95 | 0.0 |
| 20079_4 | 0.94 | 0.0 |
| 20196_18 | 0.96 | 0.0 |
| 20196_19 | 0.96 | 0.0 |
| 20239_0 | 0.93 | 0.0 |
| 20239_3 | 0.95 | 0.0 |
| 20250_1 | 0.87 | 0.0 |
| 20736_0 | 0.94 | 0.0 |
| 20944_1 | 0.79 | 0.0 |
| 21191_0 | 0.83 | 0.0 |
| 21303_0 | 0.92 | 0.0 |
| 2180_2 | 0.95 | 0.0 |
| 21817_6 | 0.87 | 0.0 |
| 2191_2 | 0.94 | 0.0 |
| 21973_9 | 0.98 | 0.0 |
| 22052_0 | 0.93 | 0.0 |
| 22091_1 | 0.78 | 0.0 |
| 2217_0 | 0.93 | 0.0 |
| 22200_0 | 0.91 | 0.0 |
| 2224_0 | 0.92 | 0.0 |
| 22408_11 | 0.97 | 0.0 |
| 22429_0 | 0.9 | 0.0 |
| 22442_11 | 0.92 | 0.0 |
| 22442_6 | 0.85 | 0.002 |
| 22475_0 | 0.9 | 0.0 |
| 2248_0 | 0.96 | 0.0 |
| 2250_0 | 0.91 | 0.0 |
| 22552_1 | 0.91 | 0.0 |
| 2256_1 | 0.98 | 0.0 |
| 22751_1 | 0.95 | 0.0 |
| 22798_0 | 0.85 | 0.0 |
| 22805_0 | 0.93 | 0.0 |

| | | |
|---|---|---|
| 22941_0 | 0.92 | 0.0 |
| 22999_3 | 1.0 | 0.017 |
| 23036_0 | 0.94 | 0.0 |
| 23279_0 | 0.96 | 0.0 |
| 23282_0 | 0.95 | 0.0 |
| 23436_0 | 0.82 | 0.0 |
| 23535_1 | 0.84 | 0.0 |
| 23593_0 | 0.95 | 0.0 |
| 23768_0 | 0.89 | 0.0 |
| 23884_0 | 0.93 | 0.0 |
| 25031_0 | 0.8 | 0.0 |
| 25084_2 | 0.9 | 0.0 |
| 25181_0 | 0.94 | 0.0 |
| 25256_20 | 0.93 | 0.002 |
| 25256_23 | 1.0 | 0.0 |
| 25284_1 | 0.9 | 0.0 |
| 25341_1 | 0.92 | 0.0 |
| 25554_1 | 0.92 | 0.0 |
| 25635_1 | 0.83 | 0.0 |
| 25818_4 | 0.94 | 0.0 |
| 25829_8 | 0.87 | 0.0 |
| 26085_4 | 0.95 | 0.0 |
| 26188_0 | 0.85 | 0.0 |
| 26212_1 | 0.91 | 0.0 |
| 26551_4 | 0.95 | 0.0 |
| 26628_4 | 0.97 | 0.0 |
| 26669_46 | 0.96 | 0.0 |
| 26988_12 | 0.79 | 0.0 |
| 26988_8 | 0.97 | 0.0 |
| 27016_1 | 0.94 | 0.0 |
| 27176_0 | 0.93 | 0.0 |
| 27689_0 | 0.91 | 0.0 |
| 28112_0 | 0.85 | 0.0 |
| 28258_0 | 0.98 | 0.0 |
| 28360_17 | 0.98 | 0.0 |
| 28360_18 | 0.96 | 0.0 |
| 28360_2 | 0.95 | 0.0 |
| 28360_30 | 0.95 | 0.0 |
| 28360_8 | 0.96 | 0.0 |
| 362_1 | 0.89 | 0.0 |
| 684_1 | 0.96 | 0.0 |
| 688_1 | 0.95 | 0.0 |
| 9936_1 | 0.97 | 0.0 |

| | | |
|---|---|---|
| 9972_0 | 0.96 | 0.0 |
| 9987_1 | 0.89 | 0.0 |

Table 5: EBG regressor correlation with the SBS ground truth of 1000 replicates. In three cases the Pearson correlation is $< 70$ (10801_0, 18850_2). For 10801_0 and 18850_2 the mean SBS support is extremely low (5 and 23) suggesting very uncertain phylogenies.
